# Quantitative Trait Loci Controlling Oil Sorption in Kenaf: Mapping and Implications

**DOI:** 10.64898/2026.03.04.709694

**Authors:** A. Emese, M. O. Balogun, R. Bhattacharjee

## Abstract

This study revealed Quantitative trait loci (QTL) controlling oil sorption capacity in kenaf which could assist in improving genotypes through marker assisted selection (MAS) for effective oil spill clean-up. Two accessions with extreme sorption capacity were selected and crossed to generate F1 which were selfed for linkage and QTL analyses. The oil sorption capacity of the F_2_ progeny was determined phenotypically and scored as a morphological marker to allow mapping of the markers associated with oil sorption gene(s). The genomic DNA extraction was carried out using a modified CTAB (cyltrimethyl-ammonium bromide) procedure. Diversity Array Technology Sequence (DArTSeq) platform was used to sequence 96 DNA samples prior to linkage and QTL analyses. Genotypic data were analysed using the JoinMap 4.1 Software to construct a genetic map of F_2_ segregating populations whereas MapQTL1 was used to detect QTL. The mapping population which included the two parents [NHC5(1) and NHC12(2)] together with 72 F_2_ progeny were genotyped with selected polymorphic SNP markers. Mapping of markers was performed using the regression mapping algorithm. 18 linkage groups were generated from the linkage analysis with Linkage group 1 (LG1) having the longest map length of 138.93 centimorgan (cM) whereas LG 16 had the shortest map length of 83.05 cM. This genetic map has a total map length of 1888.40 cM giving an average length of 104.91 cM per group and an average of one marker for every 1.39 cM. A total of 3 significant and 8 putative QTL were detected by MQM mapping and interval mapping method.

## Introduction

The exploration of crude oil and its spillage has in no doubt had serious negative impacts on our waters and agricultural lands, especially in the Niger Delta area of Nigeria, where oil spillage has been recorded at alarming levels (1). Oil spillage poses a great threat to economic development due to land degradation, loss of soil fertility, and water pollution (2). Populations of fishes in rivers and animals in forests have drastically reduced. Recent findings have shown the efficacy of using natural sorbents for oil clean-up, highlighting their importance since they do not pose environmental threats compared to synthetic sorbents and dispersants. Kenaf, an herbaceous plant, is a phytoremediant among its other beneficial properties, with strong affinity for oil.

Modified kenaf straw has been shown to absorb oil at rates nearly double that of raw straw (3), and its sorption capacity can be further enhanced through surface modification and carbonization (4). Utilizing kenaf as a natural sorbent offers a sustainable approach to managing oil spill hazards, particularly when sourced from kenaf grown in Nigeria. Additionally, its low density allows the straw to float after oil absorption, facilitating easier collection. Although kenaf is indigenous to Africa, it has not been widely utilized for oil spill clean-ups despite the high incidence of spills. This is due to insufficient awareness and the non-availability of locally adapted, high-absorbent genotypes with high yield. Consequently, improving adapted varieties will require studies on the genetic control of sorption capacity in kenaf. Reports on the genetic control of sorption capacity are, however, sparse.

Molecular breeding, which involves the use of DNA markers, is a modern approach in genetic study that provides rapid, reliable, and high-resolution results compared to conventional breeding. Examples of DNA markers include Restriction Fragment Length Polymorphism (RFLP), Random Amplified Polymorphic DNA (RAPD), Simple Sequence Repeats (SSRs), Amplified Fragment Length Polymorphism (AFLP), Single Nucleotide Polymorphism (SNP), Inter Simple Sequence Repeat (ISSR), and Diversity Array Technology Sequence (DArTSeq) (3, 5, 6). Each marker has its advantages and limitations, with some offering higher throughput and resolution than others. For instance, DArTSeq, a genotyping-by-sequencing platform, combines DArT markers with short-read sequencing to generate presence/absence and SNP markers. Its main aim is to reduce genome complexity for genotyping and has been successfully applied in kenaf for genetic mapping and diversity studies (5).

Molecular breeding assists conventional breeding through the application of molecular biological tools (7). One important tool is genetic mapping using molecular markers. In kenaf, RAPD, ISSR, SRAP, and SSR markers have been developed and successfully applied to study genetic diversity and linkage mapping (5, 8). The use of these markers has significantly facilitated the identification of traits and the development of new varieties with desirable characters at a faster and more cost-effective rate.

It is well known that most plant traits are controlled by multiple genes, and to identify such polygenic traits, quantitative trait loci (QTL) analysis is often carried out. Quantitative trait loci (QTLs) associated with key agronomic traits, including the first flowering node in kenaf, have been characterized, thereby establishing a genetic framework for the improvement of fiber yield and quality (5). However, the potential linkage between identified QTLs and sorption capacity in kenaf remains unexplored, underscoring a critical gap in current research and presenting an opportunity for integrative genetic and functional studies.

## Results

### Data filtering and construction of linkage map

DArT array returned a dataset containing 15,351 SNP markers. After data filtering was done to remove monomorphic markers and markers that were heterozygote for both parents, 3,563 SNPs were found to be polymorphic between both parents. Data analysis was prepared in the appropriate format required in Joinmap 4.1 and this was used for the linkage analysis. Segregation distortion was calculated using Joinmap 4.1 and several markers found to be distorted at a significance level p≤0.05 were removed in order to reveal the Mendelian segregation ratio as seen in Table 1. The linkage group (Table 2) formation was based on independent test LOD score done at several significance threshold level of increasing stringency ranging from 2 – 10. Mapping of markers was performed using the regression mapping algorithm. Eighteen linkage groups were generated from the linkage analysis with Linkage group 1 (LG1) having the longest map length of 138.93 centimorgan (cM) whereas LG 16 had the shortest map length of 83.05 cM as seen in Figures 4a, 4b and 4c. This genetic map has a total map length of 1888.40 cM giving an average length of 104.91 cM per group and an average of one marker for every 1.39 cM. The map comprises 1357 markers loci of SNP. Marker identification numbers are shown to the right of each LG, with map distances in centiMorgans (Haldane units) to the left.

**Table 1.**
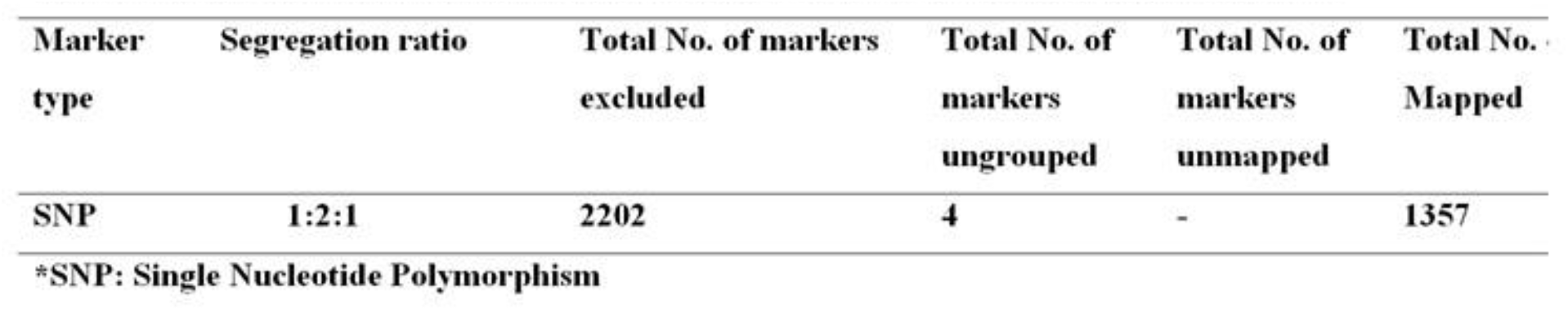
Segregation ratio and marker elimination during the construction of genetic maps.

**Table 2.**
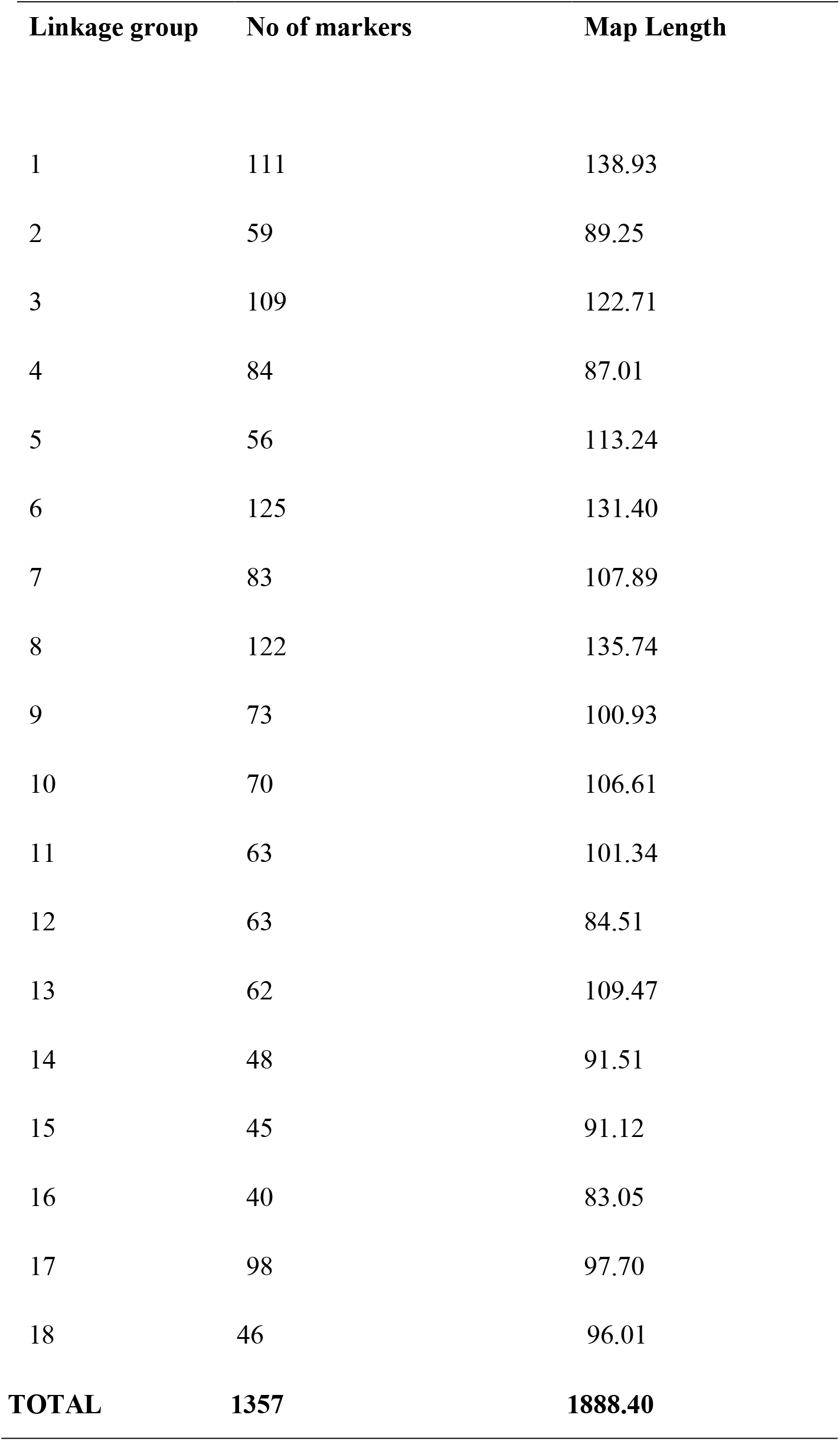
Genetic linkage groups of kenaf mapped population.

**Fig 1.**
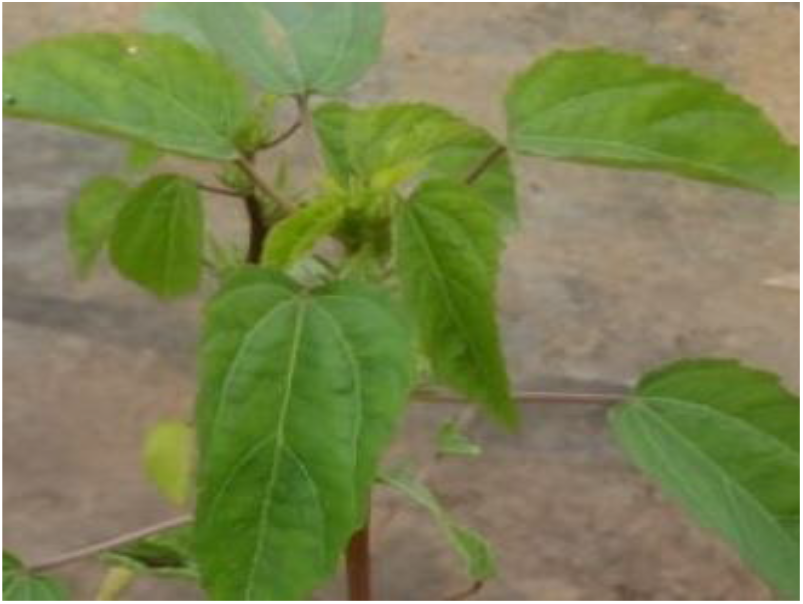
Parent 1 [NHC5(1)]

**Fig 2.**
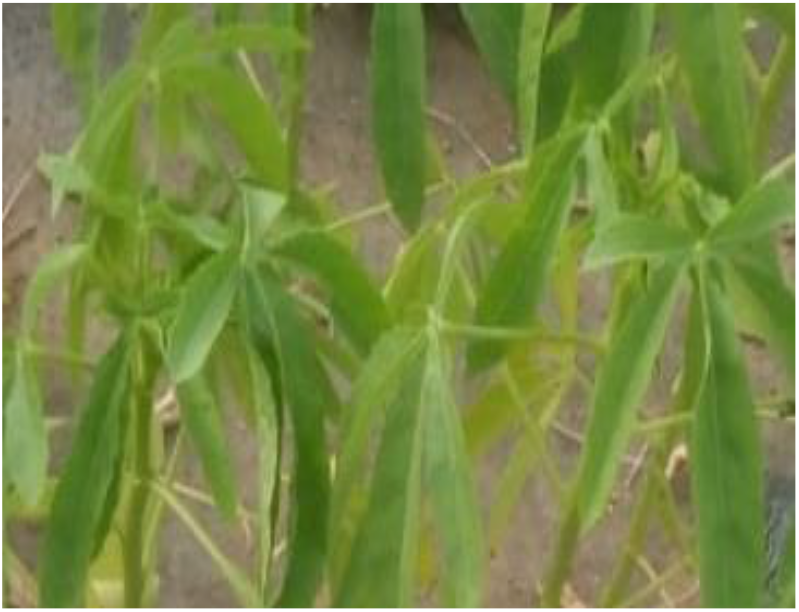
Parent 2 [NHC12(1)]

**Fig 3.**
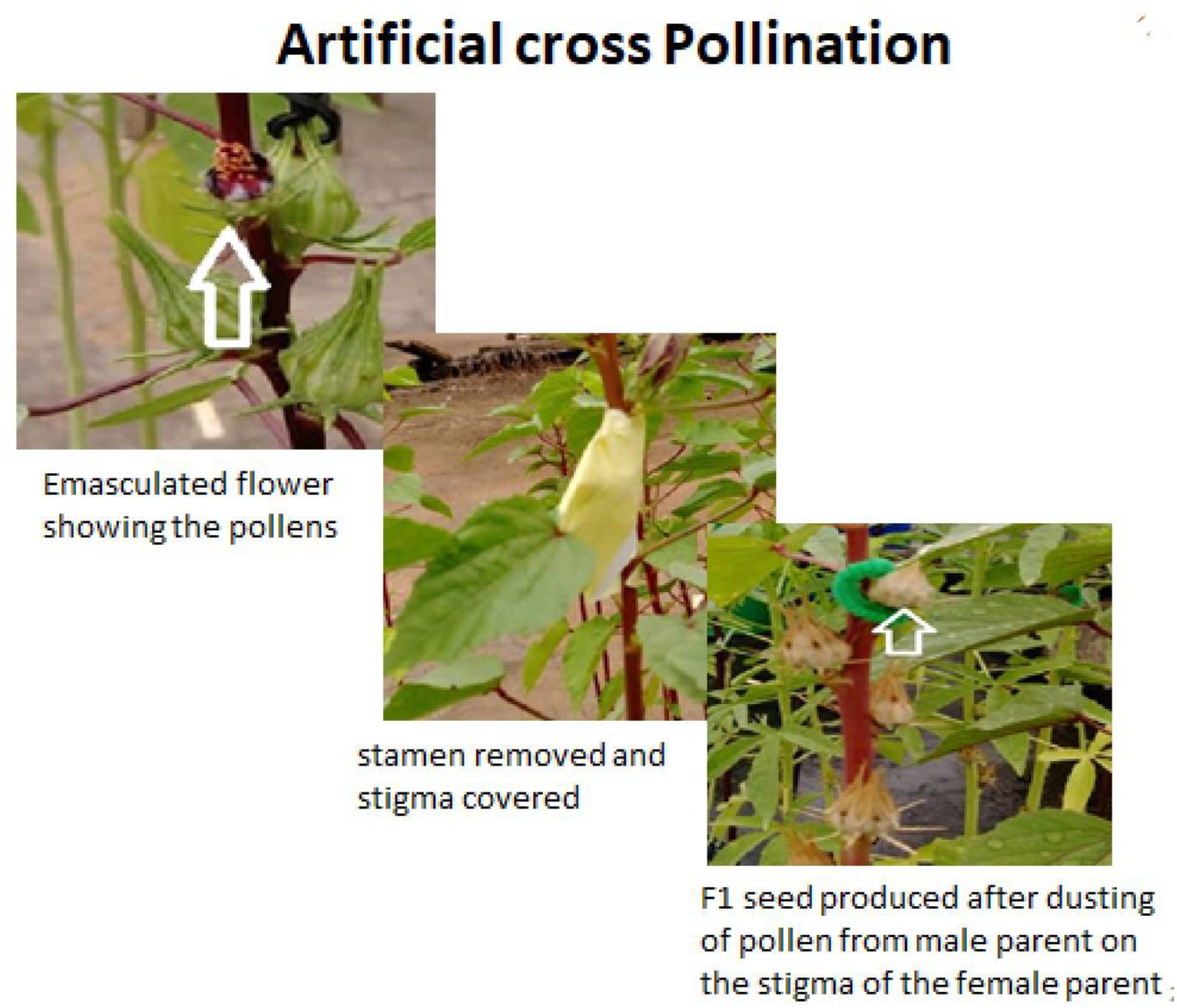
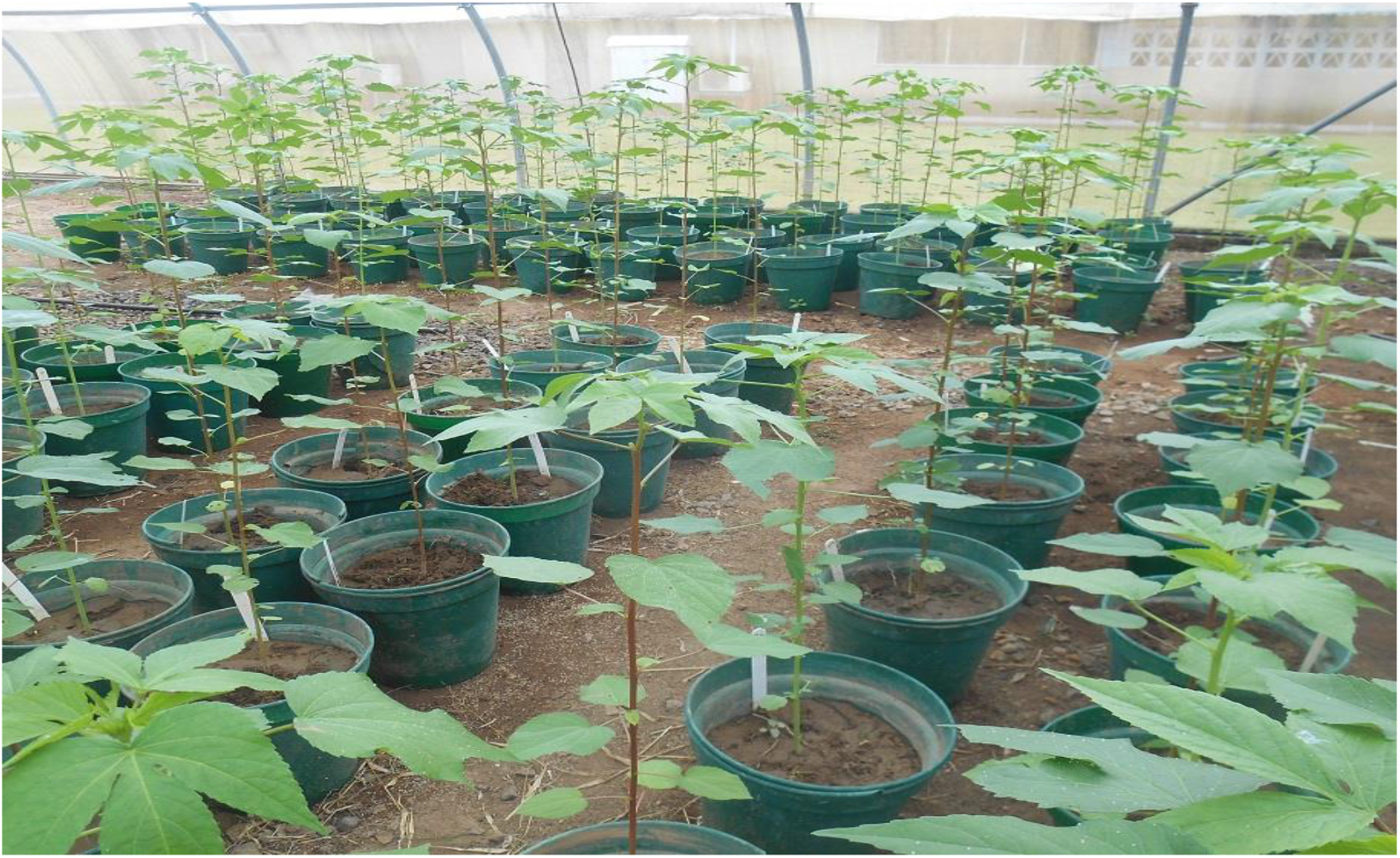
F_2_ kenaf mapping population planted at Bioscience unit, IITA, Ibadan, Nigeria.

**Fig 4a.**
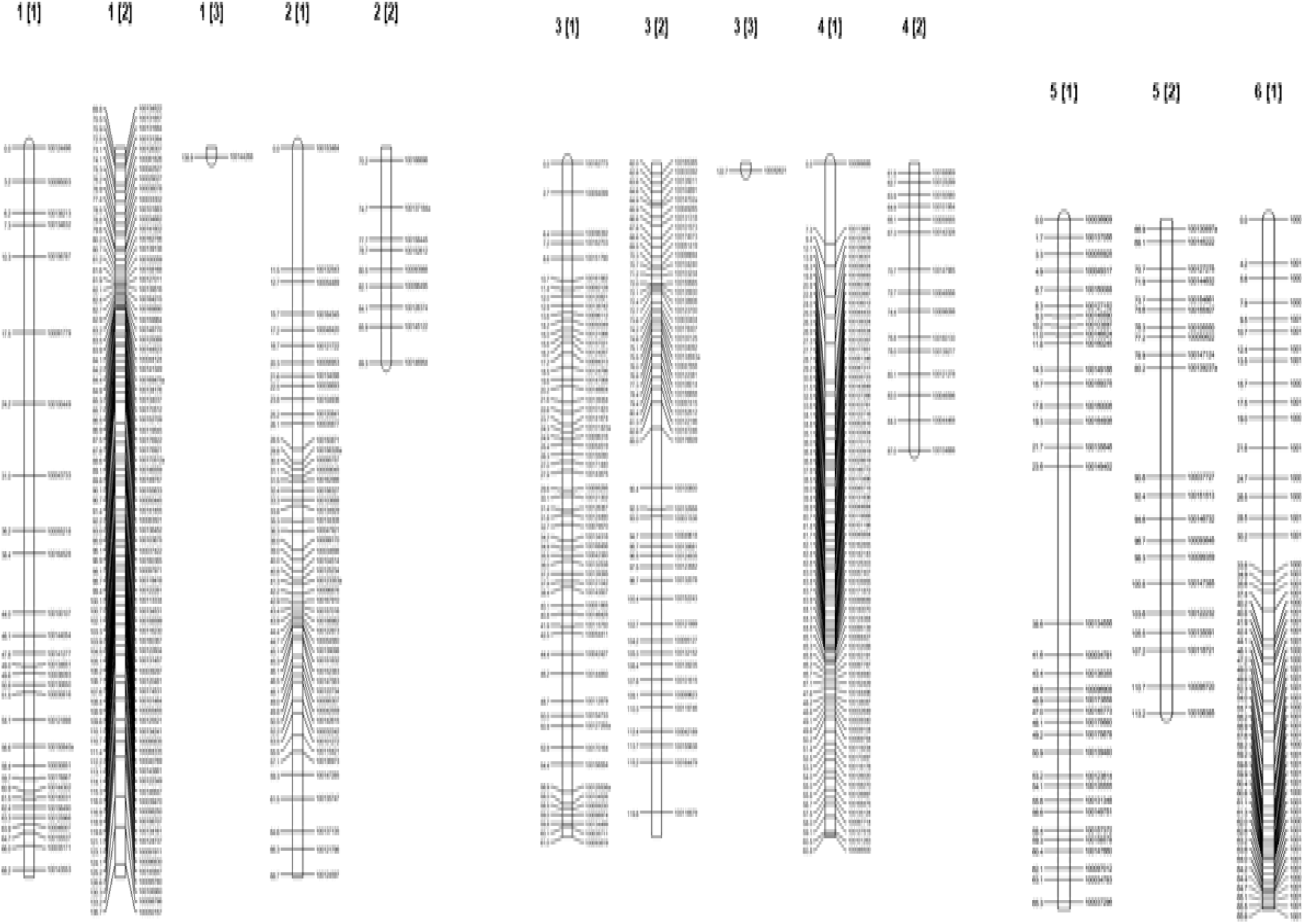
Genetic linkage map of the kenaf population showing the first two linkage groups of kenaf. Marker identification numbers are shown on the right of each linkage group (LG) with map distances in centiMorgan (Haldane units) on the left.

**Fig. 4b.**
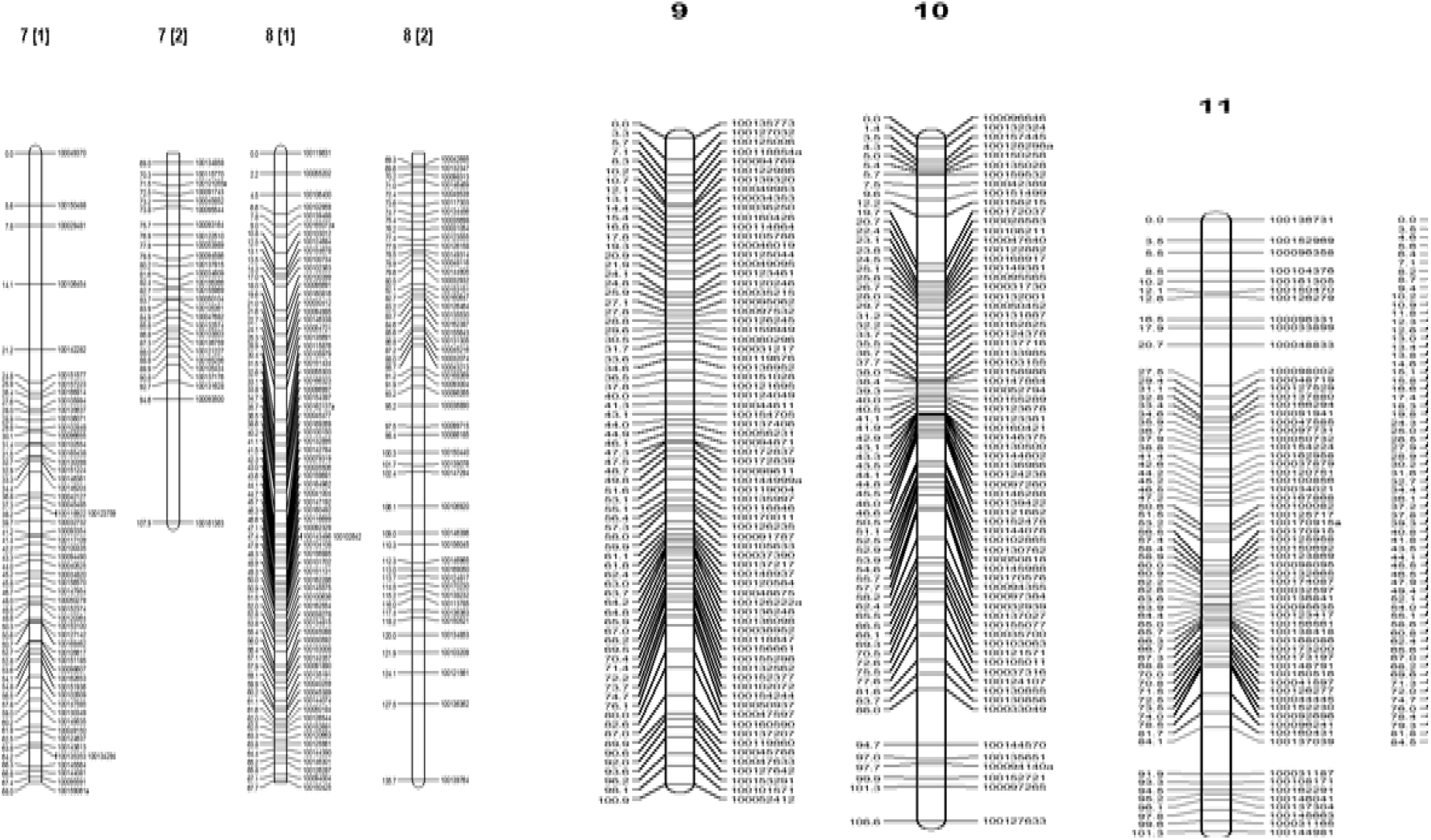
Genetic Linkage Map of the Kenaf Population Showing the Next Six Linkage Groups. Marker identification numbers are shown on the right of each linkage group (LG) with map distances in centiMorgan (Haldane units) on the left

**Fig. 4c.**
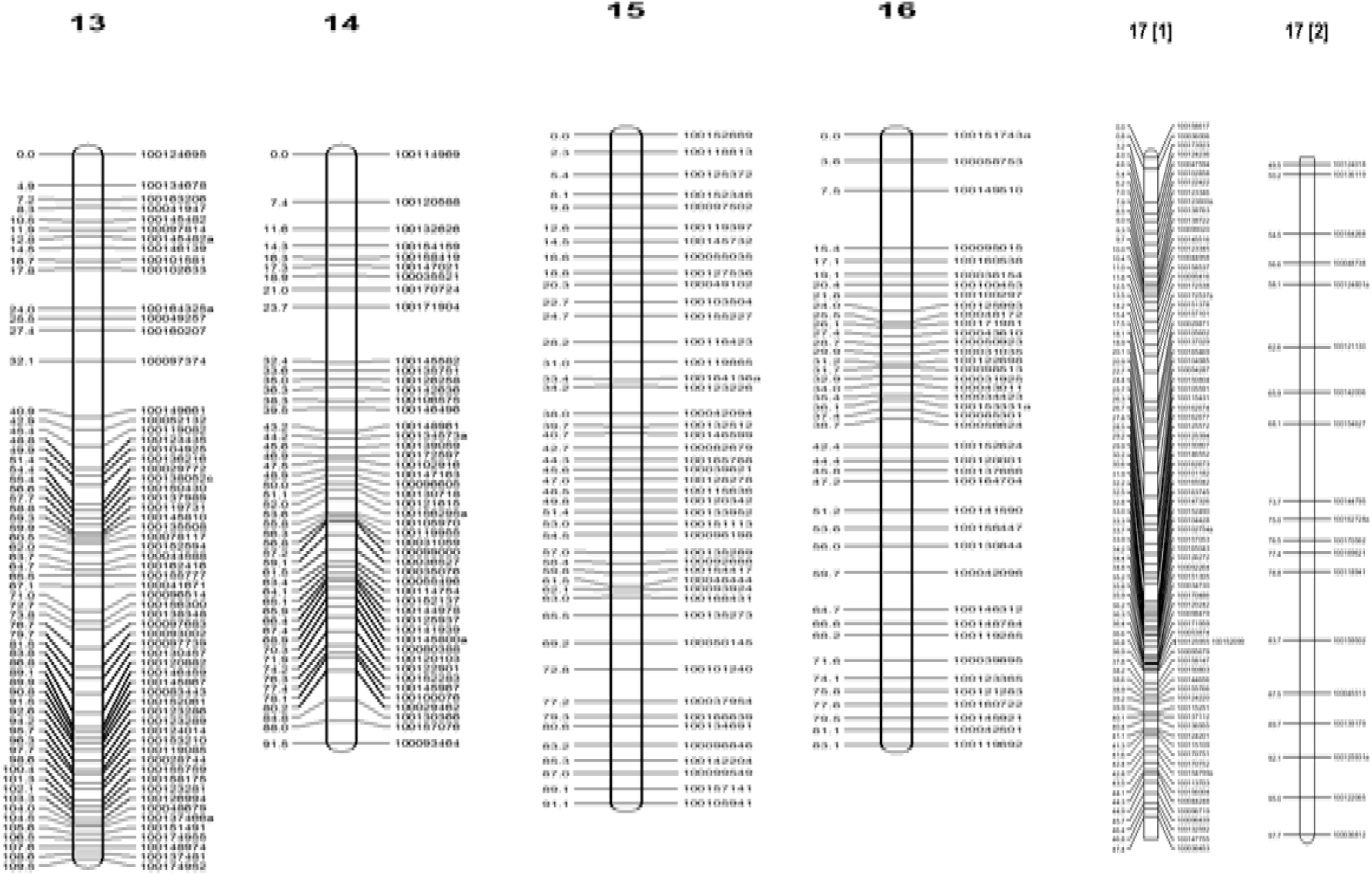
Genetic Linkage Map of the Kenaf Population Showing Eight more Linkage Groups. Marker identification numbers are shown on the right of each linkage group (LG) with map distances in centiMorgan (Haldane units) on the left.

### Identification of QTL linked to oil sorption capacity in kenaf

Significant LOD thresholds for QTL identification were determined at a genome-wide (GW) significance level of α = 5% for the sorption capacity trait using a 10,000-permutation test, which yielded a threshold value of 4.4. Using MQM and interval mapping approaches implemented in MapQTL6 software (Figs. 5a and 5b), a total of eleven QTL were identified, comprising three significant (major) QTL and eight putative (minor) QTL. Of the three major QTL, two were located on linkage group (LG) 7 at positions 24.762 cM and 36.89 cM, with corresponding LOD scores of 21.82 and 19.89, respectively. The third major QTL was identified on LG 6 at position 75.15 cM with a LOD score of 17.94 (Figure 5a; Table 3). The eight putative QTL were distributed across several linkage groups. On LG 8, three QTL were detected at positions 63.253 cM, 95.173 cM, and 62.547 cM, with LOD scores of 15.92, 13.94, and 8.42, respectively. On LG 7, two QTL were identified at positions 48.014 cM and 34.429 cM, with LOD scores of 15.5 and 5.41. Additionally, one putative QTL was detected on LG 6 at position 74.364 cM (LOD = 9.17), one on LG 12 at position 45.519 cM (LOD = 11.81), and one on LG 14 at position 32.447 cM (LOD = 11.51), as presented in Figures 5a and 5b and summarized in Table 4.

**Table 3.** Major QTLs associated with Sorption capacity identified by MQM and interval mapping.

| Trait | LG | Position | Marker ID. | LOD | Variance | % Expl. | Additive | Dominance | Cofac |
| --- | --- | --- | --- | --- | --- | --- | --- | --- | --- |
| Sorption capacity | 7 | 24.762 | 100151577 | 21.82 | 0.011461 | 16.1 | 0.898807 | 0.179996 | X |
|  | 7 | 36.813 | 100042127 | 19.89 | 0.011461 | 13.4 | -0.532587 | -0.0426269 | X |
|  | 6 | 75.15 | 100121857 | 17.94 | 0.011461 | 11 | 0.39267 | 0.104955 | X |

\*LG: Linkage group

**Figure 5a.**
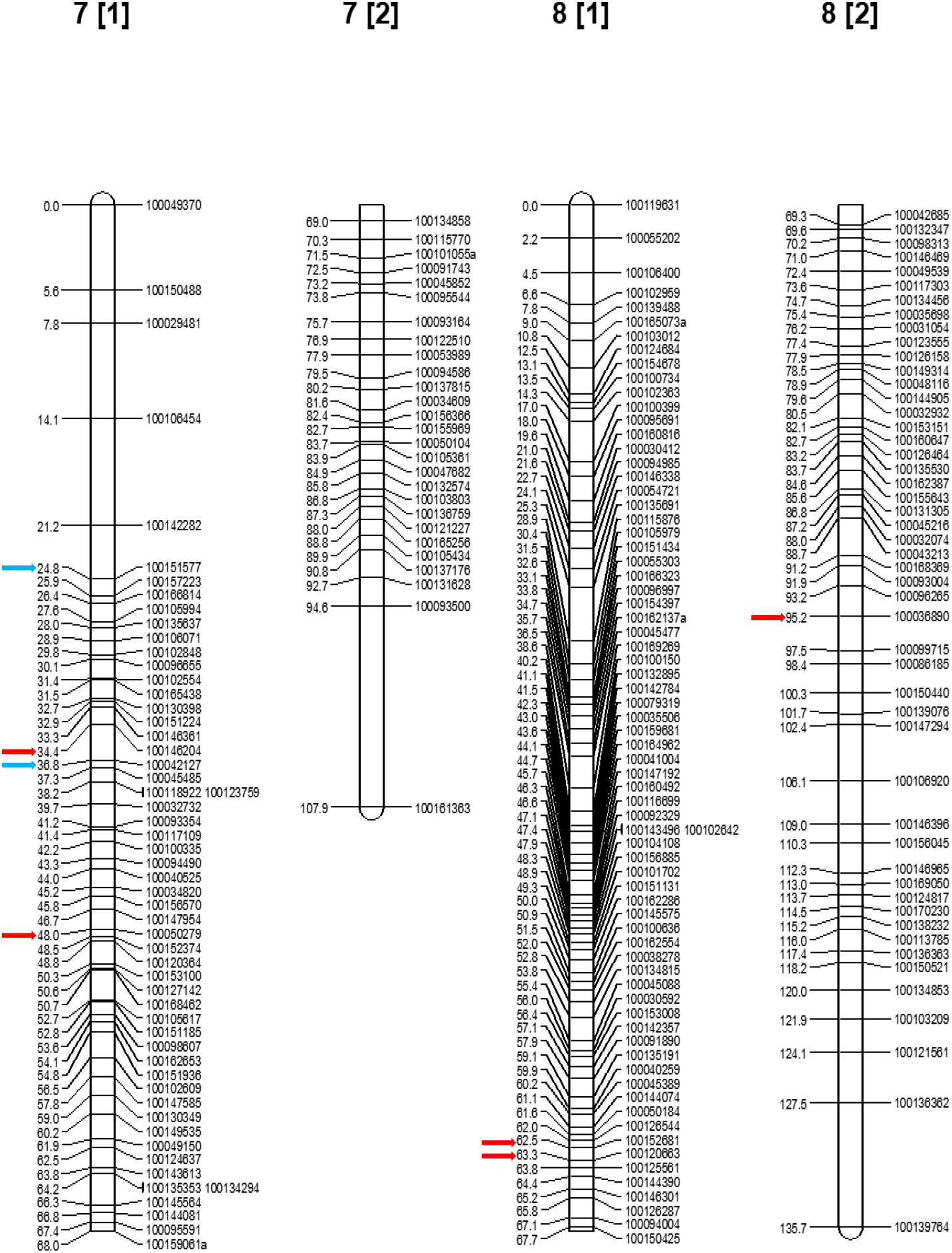
Linkage Groups Showing Quantitative Trait Loci (QTL) Linked to Oil Sorption Capacity. *Blue arrow: major QTL; *Red arrow: Minor (putative) QTL

**Figure 5b.**
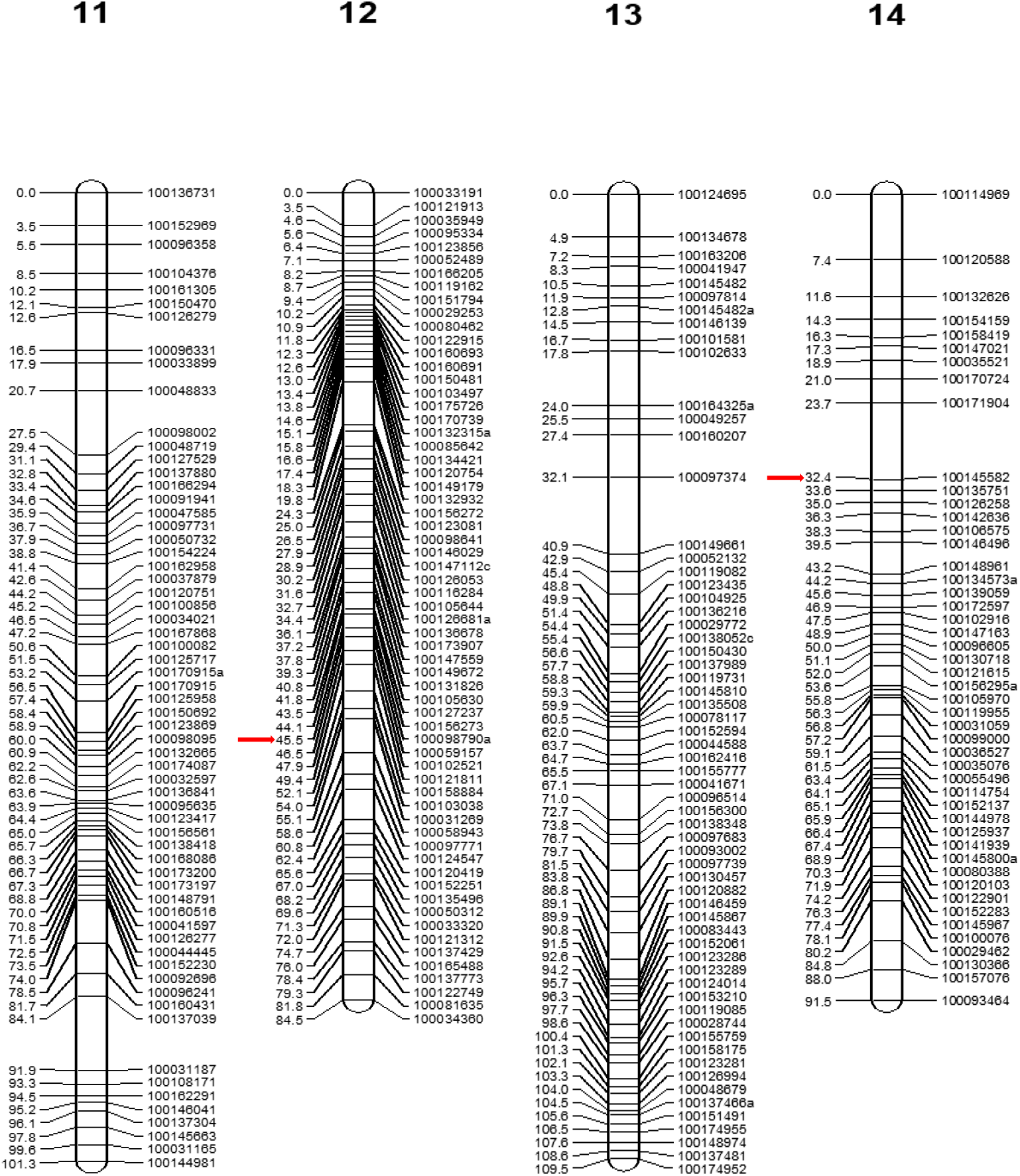
Linkage Groups Showing Quantitative Trait Loci (QTL) Linked to Oil Sorption Capacity. *Red arrow: Minor (putative) QTL

### Analysis of F2 morphological trait (sorption capacity)

Data on the sorption capacity of 72 F_2_ population were analysed to observe the distribution curve and its mendelian pattern of inheritance as shown in Figure 6. The F_2_ sorption capacity data had an overall mean of 6.43 and a standard deviation of 0.53, with mean values ranging between 4.65 g - 7.51 g.

**Fig 6.**
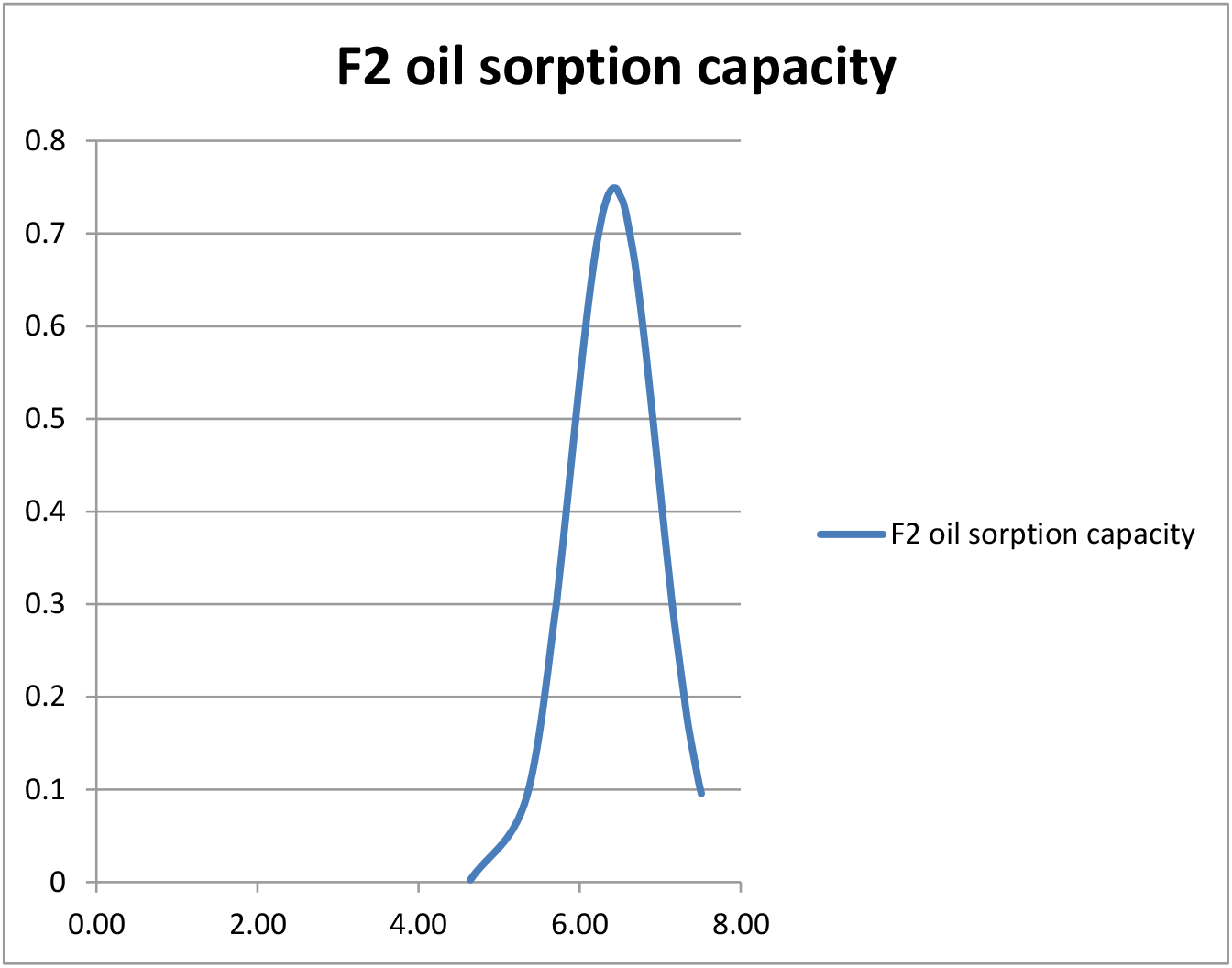
F2 morphological data for oil sorption capacity showing normal distribution curve

## Discussion

The degradation of the ecosystem as a result of continuous oil exploitation especially in Nigeria has been of major concern. Considering the use of environment friendly materials has been on the rise as efforts are being made to restore and sustain the natural ecosystem of oil-spilled affected regions. This study revealed QTL controlling oil sorption in kenaf which could assist in improving genotypes through marker assisted selection (MAS) for effective oil spill clean-up. The detection of oil sorption controlled-loci which include 3 significant and 8 putative QTL is a milestone for quick, easy and cheaper (marker assisted selection) method of genetic transformation in kenaf as regards to oil sorption trait and especially, oil sorption capacity and retention in kenaf as compared to the conventional method of pedigree, backcrossing, mass selection etc. Thus, a kenaf hybrid with good oil sorption characteristics can now be easily produced using marker assisted selection (MAS). The F_2_ oil sorption capacity data obtained revealed a normal distribution curve that show higher percentage of heterozygousity. This F_2_ segregation in accordance with Mendelian pattern of inheritance was also observed in the molecular data. Thus, both the F_2_ oil sorption morphological and molecular data confirmed the genetic diversity of the two parents [NHC5(1) and NHC12(2)] in oil sorption capacity. The findings of this study further demonstrated that the kenaf accessions utilized as parental lines in the QTL analysis are diploid, possessing 36 chromosomes. This conclusion was supported by linkage analysis, which identified 18 linkage groups corresponding to the haploid chromosome number of kenaf. It has been reported that linkage analysis ideally reflects the haploid chromosome number of the species under investigation (16, 17). If you would like, I can also make it slightly more concise or more detailed depending on whether it is for a thesis, journal article, or conference paper. Previous cytological study (18, 19) revealed all *Hibiscus cannabinus* spp forming 18 bivalents (2n = 36) at metaphase 1 and hence, are diploid. The data obtained from the F_2_ oil sorption morphological data and from the normal distribution graph showed a negative transgressive inheritance where the data obtained fell below the value obtained from NHC5(1) which serve as P_1,_ the favourable alleles but rather skewed towards NHC12(1) which served as P_2,_ the unfavourable alleles. Cytoplasmic effect must however, influenced the sorption potential of the F_2_ progenies since from the experiment, P_1_ served as the male while P_2_ served as the female.

### Recommendations

The markers linked with oil sorption capacity in kenaf are essential tools for kenaf improvement through marker assisted selection (MAS) as it remains fast and affordable when compared to conventional breeding. This discovery can possibly be applied to areas that involve the use of kenaf and kenaf related species in removing heavy metals from the soil as sorption capacity also explains the phytoremediation properties of kenaf.

### Conclusion

This study shows that the sorption capacity of kenaf is controlled by genes, and is heritable. This sorption potential is also influenced by the cytoplasmic effect of the female parents following the results from the segregating pattern of the F_2_ progenies.

## Materials and Methods

### Production of F_2_ mapping population

Two kenaf accessions grown in Nigeria with contrasting characters for oil sorption capacity were planted at International Institute of Tropical Agriculture (IITA), Ibadan, Nigeria. NHC5(1) (Fig1) which had high oil sorption capacity was crossed with NHC12(1) (Fig2) with low oil sorption capacity to produce F_1._ Hence, NHC5(1) served as parent 1 (P_1_) whereas NHC12(2) served as parent 2 (P_2_). Crossing was carried out (9). The calyx, corolla and anthers were emasculated from the flowers at 6 pm (West African Time) before they opened. Emasculated flowers were covered overnight. The flowers were opened at 7 am the following morning and the pistils were pollinated by touching the stigma with pollen grains from the stamina column of the male parent. The cross-pollinated flowers were tagged and F1 seeds were harvested at maturity. The F1 seeds were planted and flower produced on grown stems were self-pollinated to generate F2 seeds. F2 seeds were also planted to produce a mapping population (Fig. 3) for linkage and QTL analyses.

### Genomic DNA extraction procedure

The genomic DNA extraction was carried out using a modified CTAB (cyltrimethyl-ammonium bromide) procedure. 200 mg of fresh leaf samples were obtained from the two parents and their F_2_ progeny. The leaves were kept in bags under ice and immediately, lyophilized at −180°C for 48 hours pending extraction of DNA. Before extraction of DNA, hepes buffer was used to wash off secondary metabolites and mucilages in order to have quality DNA. The tubes containing the leaf samples were geno-ground and 750 ul of hepes buffer containing 2 mls of mercaptoethanol was added to each sample in extraction tubes. Samples were homogenized and centrifuged at 3500 speed (rpm) for 20 mins. The supernatant was decanted and 750 ul of hepes buffer containing 2 mls of mercaptoethanol was added to each sample thereby repeating the process. This is in order to wash off the high secondary metabolites and mucilage in the leaf samples. A solution consisting 1500 µl CTAB buffer was added to each sample in the extraction tubes. Tubes were incubated for 60 min at 55 C°, and then briefly cooled on bench for 10 min. To separate the aqueous from debris, tubes were spun for 5 mins at 1200 g. The aqueous phase (700 µl) was transferred to new tubes and 700 µl of phenol: chloroform: isoamyl alcohol (P: C: I; 25:24:1) was added. Tubes were vigorously mixed by vortex and centrifuged for 5 min at maximum speed (1300 g). Upper aqueous phase was transferred by pipetting slowly into a new tube. These two steps were repeated twice. Half volume of cold 7.5 M ammonium acetate was added. Theerafter, 2/3 volume (using the combined volume of aqueous phase and added ammomium acetate) of cold (−20°C) isopropanol was added and tubes were inverted gently. To precipitate the DNA, tubes were placed in −20°C for 30 min and centrifuged at 1300 g for 20 min to pellet the DNA. DNA pellets were rinsed with 700 µl of cold 70% ethanol and centrifuged for 5 min at 1300 g. The liquid was pippeted off and about 700 µl of cold 95% ethanol was added and centrifuged for 1 min at 1300 g. Tubes were dried and samples were re-suspended in 50 µl water by gently vortexing and then kept at −20°C.

### Determination of DNA concentration and quality confirmation

DNA concentration determination was carried out by measuring the absorbance at 260nm with spectrophotometer. The DNA concentration was calculated using the formula (DNA= optical density (OD260) x dilution factor x constant (50 ug/ml). DNA samples were diluted to a working concentration of 25ng/µl using distilled water and then stored at −20°C. Quality of DNA was examined by running it on 0.7% gel agarose for 45 min at 70 V and was later exposed to UV light. The DNA was also measured with a spectrophotometer at 260/280 readings.

### Development and characterization of DArT and SNP markers from the DArTSeq platform

The two parents and their F_2_ progeny making a total of 96 samples were sent to Diversity Arrays Technology, ILRI Campus, Nairobi in Kenya, for genotyping using DArTSeq platform. Twenty microlitres of DNA with concentration of about 50 to 100 ng/μL from each sample was pipetted into fully skirted 96 well plates (10). The plates were capped and sealed with parafilm prior to shipment for DArT services. The process of DNA genotyping using DArTSeq involved generation of genomic representations of individual samples using restriction enzymes combinations that involve *Pst*I (11). A *Pst*I-RE site specific adaptor was tagged with 96 different barcodes enabling a plate of encoded DNA samples to run within a single lane on an Illumina Genome Analyzer IIx. A sequencing primer was included in the *Pst*I adaptor so that the tags generated were always reading into the genomic fragments from the *Pst*I sites. After the sequencing run, the FASTQ files were quality filtered using a threshold of 90% confidence for at least half of the bases and with more stringent filtering for the barcode sequences. The filtered data were then split into their respective targets (genotypes) using a barcode splitting script. After producing various QC statistics and trimming of the barcode, the sequences were aligned against the reference created from the tags identified in the sequence reads generated from all the samples. The output files from the alignment generated using the Bowtie software were processed using an in-house analytical pipeline to produce a “DArT score” (presence/absence) and “SNP” tables.

### Morphological evaluation of kenaf parents and their F_2_ sorption capacity

After 150 days of growth, kenaf stems were harvested at IITA and transported to the Department of Crop Protection and Environmental Biology, University of Ibadan, Nigeria, for phenotypic evaluation. The kenaf stems were decorticated into core and bast, and air-dried. The core and bast were ground separately using a Hammer mill machine, into an average size of 0.8cm-1.2cm to form a rough core (CR) and bast (12). A combination of core rough and the bast (CBR) in a ratio of 1:1 was used to evaluate the sorption capacity of the two kenaf parents and their F_2_ progeny since it was proven to have had the highest sorption capacity as compared to other treatments. 1 g of CBR (bound with cheese cloth) was immersed in a 250 ml beaker containing 20 g of crude oil. The beaker was agitated on a laboratory shaker at 220 rpm for 5 minutes. The beaker was placed on a laboratory shaker and agitated at 220 rpm for 5 minutes. After agitation, the beakers were removed, and the kenaf treatment samples were allowed to remain suspended for 30 minutes. The sorption capacity (S), defined as the mass of oil absorbed per gram of sorbent, was subsequently determined (13) using the relationship below:

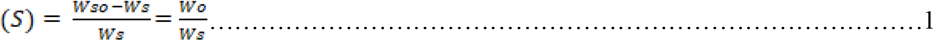

Where *W*_*s*_ is the weight of dry sorbent sample (g), *W*_*so*_ is the weight of the sorbent saturated with oil product (g), and *W*_*o*_ is the weight (g) of oil product retained in the sorbent matrix.

### Construction of genetic map and identification of QTL linked to oil sorption capacity

The mapping population which includes the two parents [NHC5(1) and NHC12(2)] together with 72 F_2_ progeny were genotyped with selected polymorphic SNP markers and the marker scores were used for the construction of the genetic maps in this study. The oil sorption capacity of the F_2_ progeny was determined phenotypically and scored as a morphological marker to allow mapping of the oil sorption gene. The JoinMap 4.1 Software (14) was used to construct the genetic maps for the F_2_ segregating populations while MapQTL6 (15) was used to detect QTL. Genome-wide LOD significance thresholds were determined independently for sorption capacity trait using the ‘Permutation Test’ function. The ‘Automatic Cofactor Selection’ tool was used interactively to identify the strongest marker cofactors on each linkage group for the sorption capacity trait. Resulting cofactors were included in the search for QTLs that exceeded the LOD significance threshold. This analysis was carried out at the Bioscience unit of IITA, Ibadan, Nigeria.

## Aknowledgements

- International Foundation for Science, Sweden, that fully sponsored this research
- Dr Nnena and Mr. Yinka for the genotypic analyses at IITA

## Authors Contribution

A. Emese: conceptualization, investigation, methodology, writing original draft, review, and editing.

M. O. Balogun: conceptualization, supervision, funding acquisition, methodology, project administration, review and editing.

R. Bhattacharjee: conceptualization, supervision, methodology, and project administration.

